# Highlighting the species diversity, pricing trends, and conservation concerns in a major ornamental fish hub

**DOI:** 10.1101/2025.02.20.639396

**Authors:** Samuel CL Ho, Jeffery CF Chan, Wing Him Lee, Yik-Hei Sung, Jia Huan Liew

**Affiliations:** School of the Environment, The University of Queensland St Lucia, QLD 4072; School of Biological Sciences, The University of Hong Kong, Pok Fu Lam, Hong Kong; Science Unit, Lingnan University, Hong Kong; School of Allied Health Sciences, University of Suffolk, IP4 1QJ, UK; School of Natural Sciences, University of Tasmania, Hobart, TAS 7005

**Keywords:** wildlife trade, exotic pet trade, freshwater ornamental fish, aquarium trade, price

## Abstract

The global ornamental fish trade presents significant challenges to conservation of biodiversity. Here we explore patterns in the freshwater ornamental fish market in Hong Kong, a major wildlife trading hub. Weekly surveys over three-months in the primary aquarium district documented 540 freshwater fish species from 73 families. We found that species occurrence in the market was largely price-driven, and minimum retail price was the strongest predictor of their frequency of occurrence. Along the occurrence-price gradient, we found that potentially undescribed species dominated the rare/expensive end of the market, potentially influenced by the anthropogenic allée effect. Alarmingly, 86 species (16%) were potentially undescribed, while 38% were either not evaluated or classified as Data Deficient by the IUCN. While common/cheaper species dominate the market, the substantial number of little-known species being sold is of conservation concern, underscoring the need for more extensive monitoring at the risk of losing them to unregulated exploitation.

## Introduction

The global wildlife trade threatens all phyla across the tree of life^1^ by increasing extinction risks^2^. On average, species abundance declines by 62% where trade occurs^3^. The persistent removal of species from their native range impacts not only the target species but can often result in ecological cascades that lead to long-term changes to ecosystem functions^4^. More than 420 million individual wild-caught animals that were listed under the Convention on International Trade in Endangered Species of Wild Fauna and Flora (CITES) were traded between 1998 and 2018^5^ as pets, sources of sustenance, status symbols, medicine, and many other purposes^1,6–8^. This is likely driven by the highly lucrative nature of the legal wildlife trade (CITES and non-CITES species) which generates an estimated revenue of USD$220 billion per year^9^. The value of the illegal wildlife trade, believed to range from US$7 billion to US$23 billion per year^10^, might also be underestimated given the proliferation of online marketplaces and transactions via social media where trade volumes can be difficult to monitor ^11,12^.

The legal status of the trade for any particular species may not reflect the long-term viability of harvesting wild populations^13^. For instance, the discovery and subsequent popularity of the boesemani rainbowfish (*Melanotaenia boesemani*) in the aquarium trade led to unregulated, albeit legal, exploitation of the species^14–16^. More than 60,000 males were caught and exported monthly^16^, causing rapid declines in wild populations^14–17^. Although this species has since been classified as “Endangered” by the IUCN, and exports are now restricted to captive-bred individuals^18,19^, the significant damage caused by the trade is likely irreversible. Worryingly, a majority of traded plants and animals are not afforded legal protections under CITES^13^. This includes hundreds of species that are considered Endangered by the International Union for Conservation of Nature (IUCN) Red-List and 93% of all described species awaiting assessment^20^. Consequently, knowledge gaps persists in the diversity and trends of the trade in unregulated and under-regulated species, which is often overlooked by large-scale studies that rely primarily on accessible databases like CITES and IUCN^1^, while underestimating the overall biodiversity impacts of wildlife trade^21^.

Inadequacies in regulations for supply-side mitigation of wildlife trade impacts suggest that there is a need to better understand demand-side drivers of the wildlife market^19,22,23^. However, we currently lack comprehensive understanding of what drives demand in the exotic pet trade^28^, including trade in the ornamental fish^29,30^. There is some evidence to suggest that price^31^, rarity and novelty underpin ownership of exotic pets^21,26,32^. Other factors such as relational dimensions (e.g companionship, ability to bond, attachment), rare aesthetic features, (smaller) body size, colors or patterns^22^ are also often sought after^28^.

Consumer preferences play a pivotal role in shaping the dynamics of the wildlife trade because this means that certain groups are more vulnerable to exploitation^23,24^. In general, consumers of the ornamental wildlife trade fall into two major groups, serious hobbyists and mass market buyers, both of whom have unique influences on market trends^23^. Overall, cheaper goods are more common in markets because they present lower barriers to entry, thus appealing to mass market buyers^25^. On the other hand, species that are rare, wild-caught, novel, or individuals that are morphologically unique (e.g., albinos), are more expensive and appeal primarily to serious hobbyists^11,21,23,26^. For the latter, the anthropogenic Allee effect^27^—in which the rarity of a species increases its demand—creates a feedback loop that may lead to extinction. This might also be the reason that CITES-listed animals and IUCN threatened animals are sometimes more expensive

Fish are a good model system for understanding consumer preferences in the ornamental wildlife market because they are the most highly traded organismal group with 15,374 species identified in the market^1^, of which >46% are traded within the ornamental market ^33^. The ornamental fish trade is also a growing sector with an estimated market value of USD$ 6.36 billion in 2023, which is likely to grow to USD$11.3 billion by 2030^34^. Freshwater fishes dominate the ornamental fish trade market, comprising approximately 5,300 species or 90% of the total trade volume^19^. Despite this, only a small fraction of freshwater fishes are protected by CITES^35^, leaving many vulnerable to unregulated exploitation. Gaining insights into the drivers behind species’ market prices and prevalence is crucial for assessing conservation risks and informing more effective regulatory measures.

In this study, we focused on the freshwater ornamental fish trade to identify patterns that might be generalizable to the wider wildlife trade. To do this, we conducted weekly quantitative market surveys in ornamental fish shops in the Hong Kong Special Administrative Region of China (hereafter shortened to Hong Kong) over a 3-month period. Hong Kong is one of the world’s largest wildlife trading hubs^6,36,37^ and a major importer of ornamental fish, with a market value of USD$19.3 million as of 2014^19^. Between 2007 and 2016, imports of exotic pets into Hong Kong increased nine-fold^36^. While ornamental fish are widely traded across Hong Kong, ‘Goldfish street’ in Mong Kok has been the largest conglomeration of pet fish stores since the 1990s^38^ and is likely a microcosm of the wider Hong Kong ornamental fish market. We undertook a comprehensive survey of ‘Goldfish Street’ by building an inventory of freshwater fish species present in the market and their respective retail prices (when available), supplementing this with information about native ranges and IUCN status. We used these to assess market trends (e.g., species popularity) and potential predictors of these trends using random forest analysis and linear regression. We then determined if there are price premiums on species considered to be threatened.

## Results and Discussion

### Overall species diversity in the trade

We recorded a total of 11,369 occurrences (i.e., observations per species per survey) comprising 540 freshwater fish species from 73 families. This is a notable increase compared to the 167 freshwater species from 50 families recorded between summer 2012 and spring 2013 in Hong Kong^39^. The diversity also surpasses figures reported by one of few published studies with a comparable study design, where authors recorded a combined total of 326 freshwater and marine species from 64 families from a year-long survey across Greece^31^. Of the 540 species encountered, 86 lacked formal scientific description yet were recognized as a distinct “species” in the market. We cataloged these at the genus level, along with unique identifiers (e.g., *Panaqolus* sp. L397). They were either undescribed species or variants marketed as distinct species, which we collectively grouped into a category named undescribed (UN) for subsequent analysis. Additionally, two “species” (i.e., *Amphilophus citrinellus* × *Vieja melanurus* and *A. citrinellus* x *Paraneetroplus synspilus*) were hybrids.

We used species accumulation curves to estimate the completeness of our surveys and found that expected species richness approaches saturation after our 14 surveys (Supplementary Figure 1). The extrapolated total species richness within the market was 585 species (i.e., average output from Chao, First order Jackknife, Second order Jackknife and Bootstrap models), or 476 with a more conservative approach that included only 453 formally described species from our surveys. . While these estimates exceed those of any comparable surveys, they are still likely to be an underestimate of the overall diversity in the market because we omitted a small subset of shops that were too logistically challenging to survey consistently due to unpredictable opening hours and lack of accessibility (e.g., located in upper levels of mixed-used building complexes). These often include specialty shops for cichlids, armoured catfish, and corydoras, among other species that are sold rarely in non-specialized outfits.

Among the 73 families observed, Loricariidae, Charicidae, Cichlidae, Callichthyidae and Cyprinidae (in descending order) dominated the market in terms of the number of species in the market. This is largely consistent with the findings from 2012–2013 market surveys in Hong Kong, where Cyprinidae and Cichlidae, Osphronemidae, Poecilidae and Characidae (in descending order) were the most common families in the trade^39^. Other families were represented more sporadically. Our data show that 59 of the families encountered were represented by five or less species in the trade, while 32 families were represented by only one species.

The majority of species and families in the trade are native to South America and Southeast Asia (Fig. 1). Only two percent of fishes (13 species) were native to East Asia, including four species being native to Hong Kong: Chinese barb (*Barbodes semifasciolatus*), blue neon goby (*Stiphodon atropurpureus*), Hong Kong paradise fish (*Macropodus hongkongensis*), and fork-tailed paradise fish (*Macropodus opercularis*)^40^. These trends might be explained by the rich diversity of freshwater fishes in Brazil^41^ and Southeast Asia in general. Moreover, Brazil and Singapore are among the world’s largest exporter of freshwater ornamental fish^42^.

**Fig. 1:**
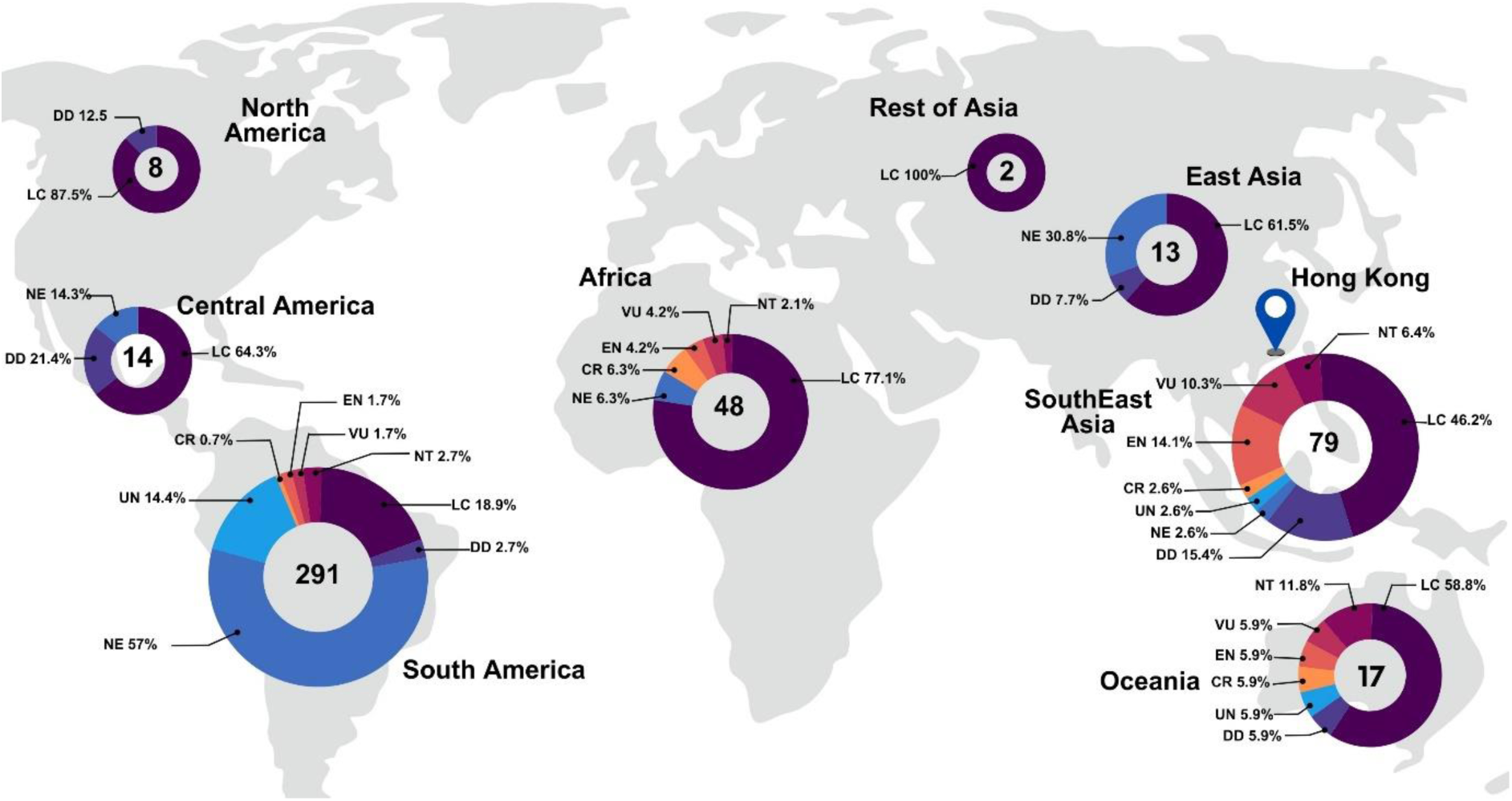
Continents and subcontinents where fishes from the market are native to. The pie chart indicates the proportion of IUCN statuses of fishes from the region. The number within the pie chart indicates the number of fishes sold from that native origin.

### Market trends

We quantified a species’ market occurrence by calculating its frequency of occurrence per shop, per survey. We used a Random Forest machine-learning approach to identify possible drivers of market occurrence in trade and found that minimum retail price to be the most useful predictor. Upon further examination using linear regression we found that lower-priced species were more likely to be traded (Estimate = -0.0010722, 95% CI [-0.001416, -0.000728], p < 0.001). Other price-related predictors, including average price, price range, max price, and price change were more useful in explaining patterns in species occurrence than non-price-related variables including IUCN status, family, and native origin (Fig. 1).

Having established a clear relationship between market occurrence and price, we assessed how each species placed within the market occurrence-minimum retail price gradient (Fig. 2), which could be indicative of whether the species appeals to mass-market consumers (common/cheap) or serious hobbyists (rare/expensive). Nine of the 10 species with the highest minimum price were from the Loricariidae family (Fig. 2) native to South America. Loricariids were encountered less frequently than most other fishes in our surveys, with a mean market occurrence of 9.8, compared to the overall average of 16.2. Species in the rare/expensive end of the market are likely to be of greater conservation concern, given the likely influence of the anthropogenic Allee effect on this segment of the market^27^.

**Fig. 2:**
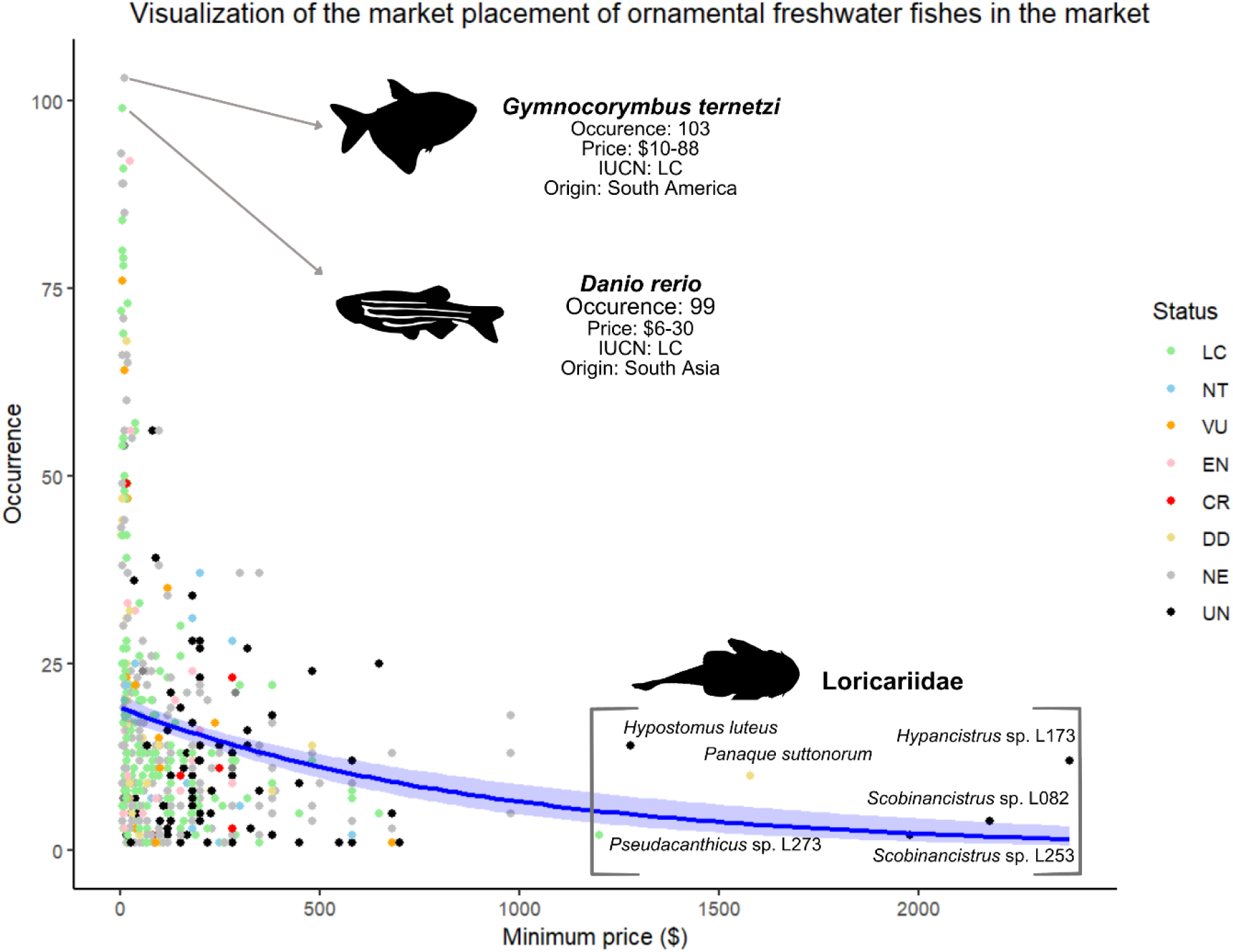
A visualization of the market placement of every ornamental freshwater fish species recorded in the market. Each dot represents a unique species, colored by their IUCN status. The blue curve and shaded blue area represent the negative correlation between the minimum price and occurrence of fishes being sold and the 95% confidence interval respectively (Estimate = -0.0010722, 95% CI [-0.001416, -0.000728], p < 0.001).

At the common/cheap end of the market appealing to mass market consumers, the black tetra (*Gymnocorybus ternetzi*) and zebra fish (*Danio rerio*) emerged as the most frequently occurring species in our survey. Both are among the cheapest in the market with minimum prices of US$0.8 and US$1.3, respectively (Fig. 2). The guppy (*Poecilla reticulata*) and neon tetra (*Paracheirodon innesi*) are the most commonly traded species across the world, and unsurprisingly ranked third and fifth most common in our study^43^. Both species were again among the cheapest species on the market, selling at US$0.3 (guppy) and $1.3 (neon tetra), respectively. Species in this segment of the market are generally captive bred at scale^49^, and are therefore of less conservation concern.

In some exotic pet markets, CITES and IUCN status confers price premiums, and species of conservation concern are often more expensive^46^. We assessed this within the freshwater fish market, and found that out of the 482 described species, 186 have not been evaluated (NE) by the IUCN. The remaining 177 were classified as Least Concern (LC), 18 Near Threatened (NT), 19 Vulnerable (VU), 20 Endangered (EN), 8 Critically Endangered (CR) and 26 Data Deficient (DD). Nineteen percent of the species encountered in the market lacked formal assessment of their taxonomic identity or conservation status, which we placed into a new category named “Undescribed” (UN). The UN category comprises mostly of members of Loricariidae, Callichthyidae, both groups are extremely diverse with hidden richness^50^. Newly imported members that were scientifically unidentified were given code names, each code usually represents a unique morphological form or locality of a species, which potentially represents an undescribed species^51^. Notably, South America had the highest percent of non-evaluated (NE; 56.9%) and undescribed (UN; 14.5%) species, contributed largely by Characidae, and Loricariidae respectively.

We then investigated the relationship between conservation statuses and retail price using the Kruskal Wallis Test followed by the Post-hoc Dunn’s Test^54^. We found that UN species were priced significantly higher than LC (*Z = -7.00, p < 0.001, Bonferroni-adjusted*), VU (*Z = 3.80, p = 0.001, Bonferroni-adjusted*), EN (*Z = -3.91, p < 0.001, Bonferroni-adjusted*), DD (*Z = -4.62.00, p < 0.001, Bonferroni-adjusted*) and NE (*Z = -5.48, p < 0.001, Bonferroni-adjusted*) (Fig. 3). UN species in this study were usually associated with unique morphologies, new source locations, or are potentially be undescribed ^50,51^, so our findings suggest that consumers of ornamental trade might be motivated by novelty and/or rarity. This aligns with similar observations in other wildlife markets such as the reptile and amphibian trade^21,26^.

**Fig. 3:**
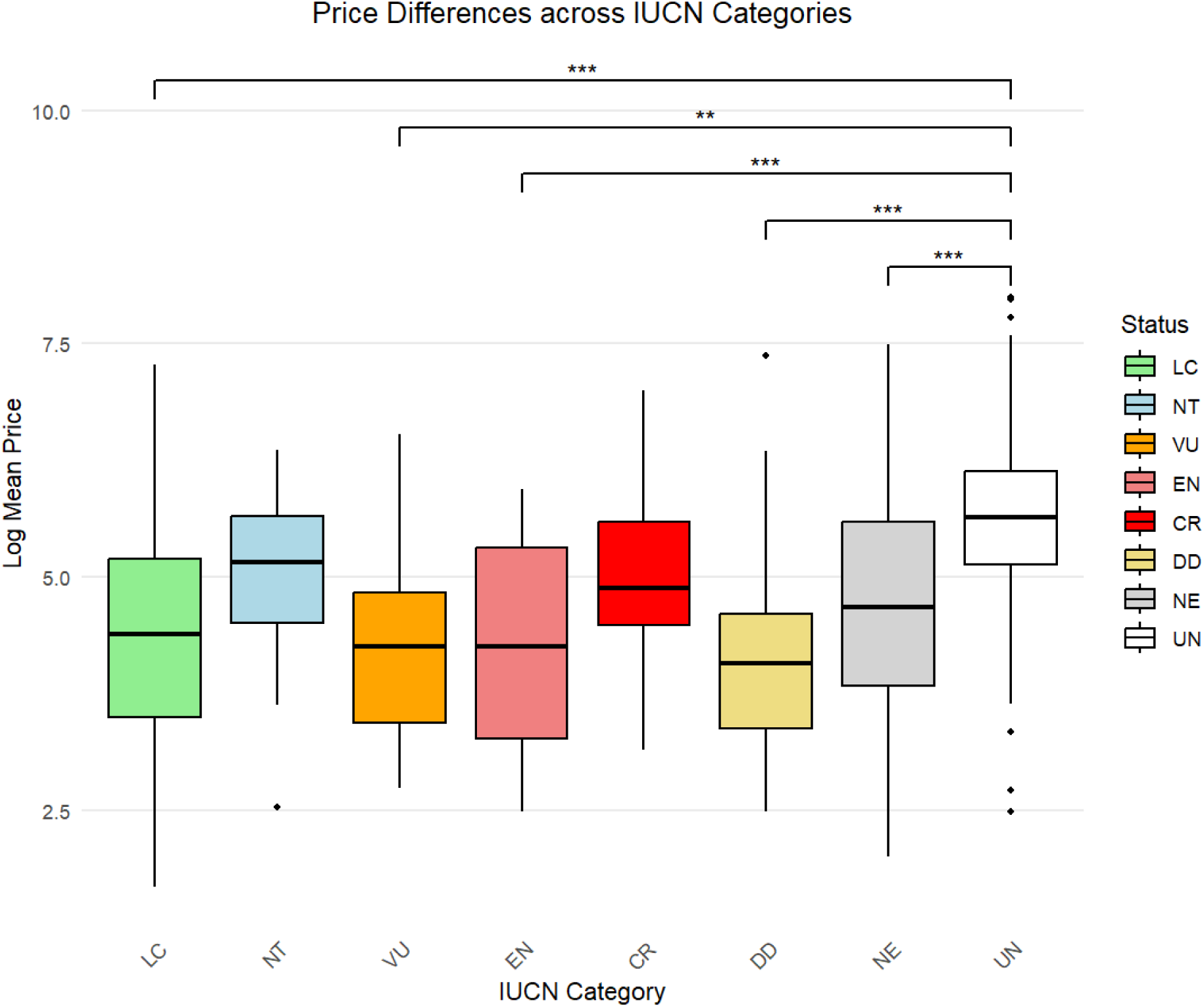
The log-transformed mean price differences across different conservation statuses, including Least Concern (LC), Near Threatened (NT), Vulnerable (VU), Endangered (EN), Critically Endangered (CR), Data Deficient (DD), Not Evaluated (NE), and Undescribed (UN). UN species had significantly higher than LC (*Z = -7.00, p < 0.001, Bonferroni-adjusted*), VU (*Z = 3.80, p = 0.004, Bonferroni-adjusted*), EN (*Z = -3.91, p < 0.001, Bonferroni-adjusted*), DD (*Z = -4.62.00, p < 0.001, Bonferroni-adjusted*) and NE (*Z = -5.48, p < 0.001, Bonferroni-adjusted*).

### Conservation implications

The trade of species that lack formal descriptions or conservation assessments poses significant risks to conservation^13^. Without reliable information on their taxonomy or population status, these species may be vulnerable to overexploitation before conservation measures can be implemented^13^. This is not without precedent, however. A newly discovered species of pencil fish, known as *Nannostomus* sp. “Super Red Cenepa”^52^, was first sold in Hong Kong pet stores within a month of its discovery, ahead of any other ornamental fish markets^52^. Other examples of widely traded but poorly understood groups include the snakeheads (*Channa* spp.), which have been subject to a six-fold increase in exported wild-caught populations from 2014 to 2019 to major markets, including Hong Kong^53^.

Besides undescribed species, we recorded several species of notable conservation concern in our survey that were known to be threatened. For example, the red line torpedo barb (*Puntius denisonii*) from the Western Ghats in India was the species with the fifth highest occurrence. The species was first traded in 1996, but its popularity in the ornamental fish market led to over exploitation in its native range^55^. Consequently, the IUCN threat status of the species was raised to “Vulnerable” in 2009, and again to “Endangered” in 2015^56^. Other examples of endangered species in the market includes the wild zebra pleco (*Hypancistrus zebra*) and the clown loach (*Chromobotia micracanthus)*, both of which are threatened by over-exploited in their native range^19^.

Most fishes (78.5%) we recorded originate from the tropics; however, we could not identify the source populations of fishes encountered in our surveys. While 90% (in volume) of freshwater species are captive-bred^33^, a significant proportion (∼10%) are thought to be wild-caught, encompassing of a wide variety of species^33^. Details concerning the source stock of animals in the trade are disclosed on CITES databases, but only a small fraction of freshwater species are covered by the convention^35^. These are primarily large charismatic species such as stingrays, sturgeons, and paddlefish^35^. The only confirmed sighting of a CITES-listed species in our market survey is the Asian arowana (*Scleropages formosus*), with retail prices ranging from US$231–5,104.

Several species recorded in our surveys are likely to be sourced from local wild populations. For example, the Hong Kong paradise fish (*Macropodus hongkongensis*) is sparsely distributed in Hong Kong, the Guangdong Province, and the Hainan Province in south China^57^, and although there is no direct evidence that individuals in the market are wild caught, the species had historically been sourced from the wild^58^. In addition, diadromous fishes, such as species from the subfamily Sicydiinae, cannot be captive-bred due to their amphidromous life cycle. Therefore, the nine Sicydiine goby species we found sold in stores were almost certainly wild-caught. This includes the cyclops goby (*Lentipes dimetrodon*), a native of Indonesia currently assessed to be “Vulnerable” ^59^. The cyclops goby was recorded once from a single shop across our entire survey period. The species was no longer detected in our surveys a month after it was recorded, potentially indicating a high turnover rate and possibly low supply as they were not subsequently replaced.

Despite the obvious impacts on biodiversity, the ornamental fish trade still divides opinions because of potential contributions to local economies and conservation efforts^68^. The ornamental fish trade can support local communities by creating job opportunities and incentivizing environmental stewardship if well-managed^19,33,68,69^. The presence of wild-caught specimens in the trade had also facilitated the documentation of the life-history traits of rare or understudied species in the past^70^. Niche hobbyist who purchase these specimens occasionally publish their observational and experimental findings on fish behavior, ecology and biology in. various forms^70^ while contributing to captive-breeding efforts. Successful commercial breeding could also play a role in breaking the feedback loop of the anthropogenic allee effect^71,72^. In addition, critically endangered species such as the red-tailed blue shark (*Epalzeorhynchos bicolor*), can sometimes be brought back from near extinction by successful commercial breeding^19,73^. Similarly, other “success” stories among species encountered during our surveys include the zebra pleco (*Hypancistrus zebra*) and the Boesemani rainbowfish (*M. boesmani*). Nevertheless, we note that these “successes” are greatly outnumbered by species that are negatively impacted, where most species were at risk due to exploitation by the pet trade in the first place^73^.

We propose that the scope of CITES should be expanded to include a broader range of threatened freshwater fishes, particularly endangered species that are known to be actively sourced from wild populations. For instance, exporters should be made to adopt an organized coding system including species name, source (wild-caught or captive), source of collection, and name of collector^33,60^. Conversely, importers should be made to adopt mandatory origin labeling, requiring traders to disclose whether species are wild-caught or captive-bred. Hong Kong as an international trading hub is well placed to take a leading role by implementing origin disclosure laws and provide incentives for the trade of sustainably sourced or captive-bred species.

### Conclusions

Our study presents a snapshot of species diversity and market trends of freshwater ornamental fishes available in Hong Kong’s ornamental fish trade. We found a strong correlation between species occurrence/rarity and price, not unlike other commodity markets. We also identified key species in the mass-market and specialized hobbyists segment of the market which includes potentially undescribed species. The unprecedented diversity of fishes found in our surveys (given the size of the market) highlights the potential threats to wild populations of freshwater fish. Most species in the Hong Kong market are yet to be evaluated by the IUCN, indicating significant gaps in our knowledge of these species. Our findings highlight the urgent need for information about species in the trade to inform sustainable practices and improved regulatory measures to strike a better balance between biodiversity conservation and economic benefits.

## Materials and Methods

### Market Surveys

Surveys were conducted weekly on Sundays in Goldfish Street (Tung Choi Street, Mong Kok, Hong Kong) from 27^th^ November 2022 to 26^th^ February 2023. To establish a comprehensive baseline of the market’s species diversity, an initial market-wide survey encompassing all 34 ground-level shops was conducted. During the baseline survey, we recorded the name of each species (as labelled in the store) and the corresponding price per individual. For guppies (*P. reticulata*) where shops displayed multiple morphs and variants of the same species, we recorded the species name once to avoid duplication. Following the market-wide survey, we selected 10 shops for repeat surveys. The 10 shops were selected based on the diversity of species being sold, with the selected shops collectively accounting for more than 90% of the total species diversity observed during our baseline survey. This approach ensured that the repeated surveys were representative of the overall market trends while focusing on a manageable subset of shops.

After each survey, we matched the labelled name to the corresponding species by cross-checking against journals, books, authentic online databases and other published documents ^76^. In cases where discrepancies arose between sources where we were unable to confidently identify the species, we omitted those observations. Ambiguous species names that could not be conclusively matched were also omitted. For species from the families Loricariidae and Callichthyidae, where some species were identified by codes (e.g., *Panaqolus* sp. L397), our primary source of information come from Novák et al.^51^, planetcatfish.com, and Corydorasworld.com. When sources suggest that the labelled name refers to a potentially undescribed species, we left them at genus level plus a unique identifier (usually their distinct code name). All verified species names follow Eschmeyer’s Catalog of Fishes^77^.

For each species, we also collated data of their IUCN red list status and their native geographic distribution from IUCN Red List of Threatened Species^78^ (accessed in February 2023). For undescribed Loricariids and Callicthyids that were assigned unique identifiers, we obtained their native geographic distribution from planetcatfish.com, and Corydorasworld.com.

We quantified a species’ market occurrence by using its frequency of occurrence per shop, per survey. Therefore, if a fish species is observed in multiple tanks of a single shop during a single survey, market occurrence of the species is counted once. This method of quantifying occurrence minimizes the effects of inflation of occurrence by shops that specialize in selling different variants of the same species of fish.

### Statistical Analyses

To assess whether our surveys had reached market saturation, we generated species accumulation curves using the “specaccum” and “plot” functions from the vegan package^79^. These curves were used to assess the cumulative number of species recorded over the course of the survey. Subsequently, we extrapolated the species richness upon achieving saturation using the “specpool” function. We obtained our final projected number of species by using the averaged estimates from the four models “specpoool” (Chao, First order jackknife, Second order Jackknife and Bootstrap). This was conducted in two parts: first, with all fish species, including those that have yet to be formally described; and second, with a smaller pool of described species to obtain a more conservative estimate of species diversity in the market. We removed shop number “20” from the analysis as it was closed on 2 separate occasions during our market surveys.

We had multiple price variables for each fish species, including its average price throughout the survey, minimum price (the lowest available price of a species in the market), maximum price (the maximum available price of a species in the market), price range (the difference between maximum and minimum price), and price change (price difference between the first and last occurrence of a species). These price variables were derived from the price data collected from shop labels throughout the entire survey period. Using all the price variables, IUCN status, and native origin, we examined the importance of each variable in explaining fish occurrences using random forest analysis with the package *ranger*^80^ . The model was trained to predict the occurrence of species from our variables with the default settings of using 10,000 trees and permutation importance. We modelled the most representative variable from random forest analysis (minimum price) with “Market Occurrence” using a negative binomial generalized linear model (*glm.nb* function in R package *MASS*)^81^

We assessed the price variability between IUCN categories using the log-transformed mean price of each species. We tested for normality of our data using the Shapiro-Wilk test using the function *shapiro.test* in R package *stats* and proceeded to investigate whether log mean price varies significantly between categories with the non-parametric Kruskal-Wallis Test using the function *kruskal.test* in R package *stats*. The differences between each group were further tested using post-hoc Dunn’s test using the function *dunnTest* in R package *FSA*. We tested the homogeneity of variances using the Leven’s test and Bartlett’s test from the R packages *car* and *stats* respectively. .All data processing and analyses were conducted in the R environment^54^.

## Data Availability

Primary data generated and analyzed in this study is available through figshare: https://figshare.com/s/54f661837e729cd8508a, DOI: 10.6084/m9.figshare.27001738. All other data is available from the corresponding author on reasonable request.

## Statements and Declaration

The authors have no relevant financial or non-financial interests to disclose. The authors declare no competing interests.

## Author Contribution

All authors contributed to the study conception and design. Data collection was performed by SCLH and WHL. Analysis was performed by SCLH and JHL. The first draft of the manuscript was written by SCLH and all authors commented on previous versions of the manuscript. All authors read and approved of the final manuscript.

## Acknowledgements

We would like to acknowledge the Freshwater Collective for their support throughout the study. We thank Lauren Ohayon and Aditya Pramudya for providing feedback on the manuscript.

## Supplementary Material (also provided in a separate document)

**Fig. 1:**
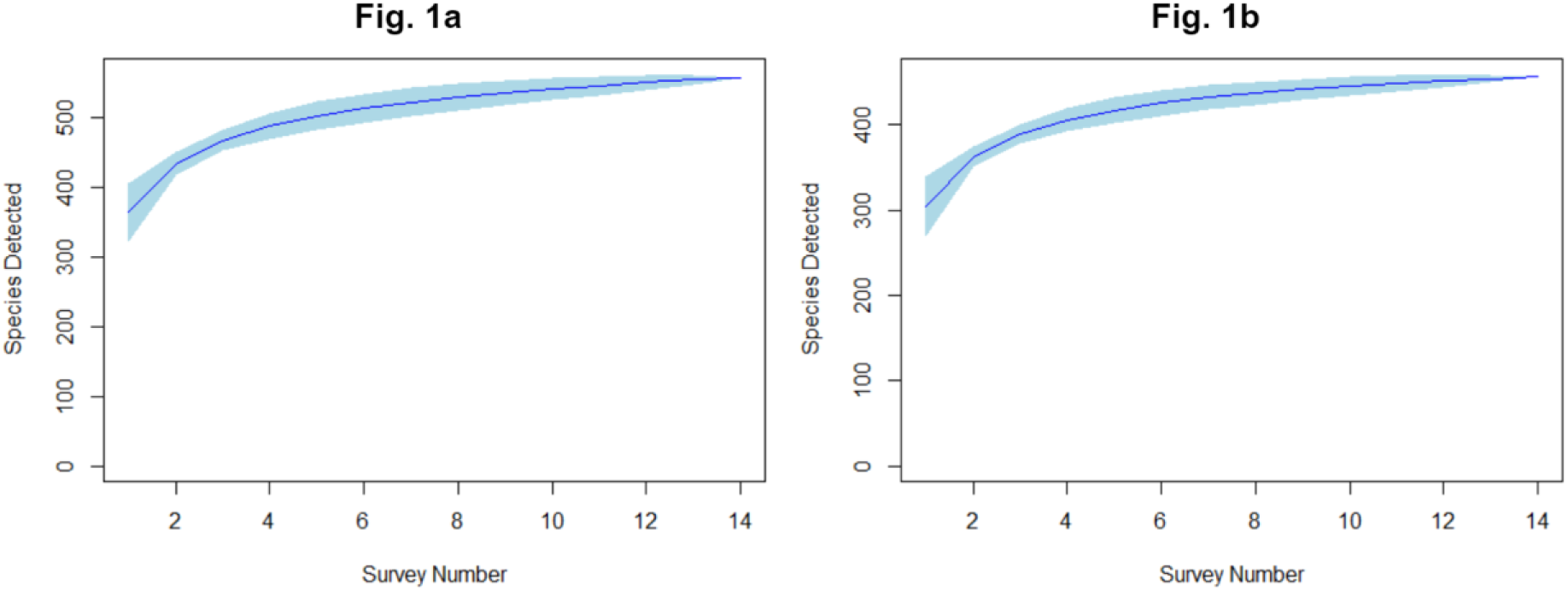
species accumulation curves from 14 weekly surveys in Gold Fish Street, Mong Kok, Hong Kong using (a) all fishes, and (b) only scientifically recognized fishes. The continuous lines represent the mean and the shaded areas the 95% confidence interval.

**Fig. 2:**
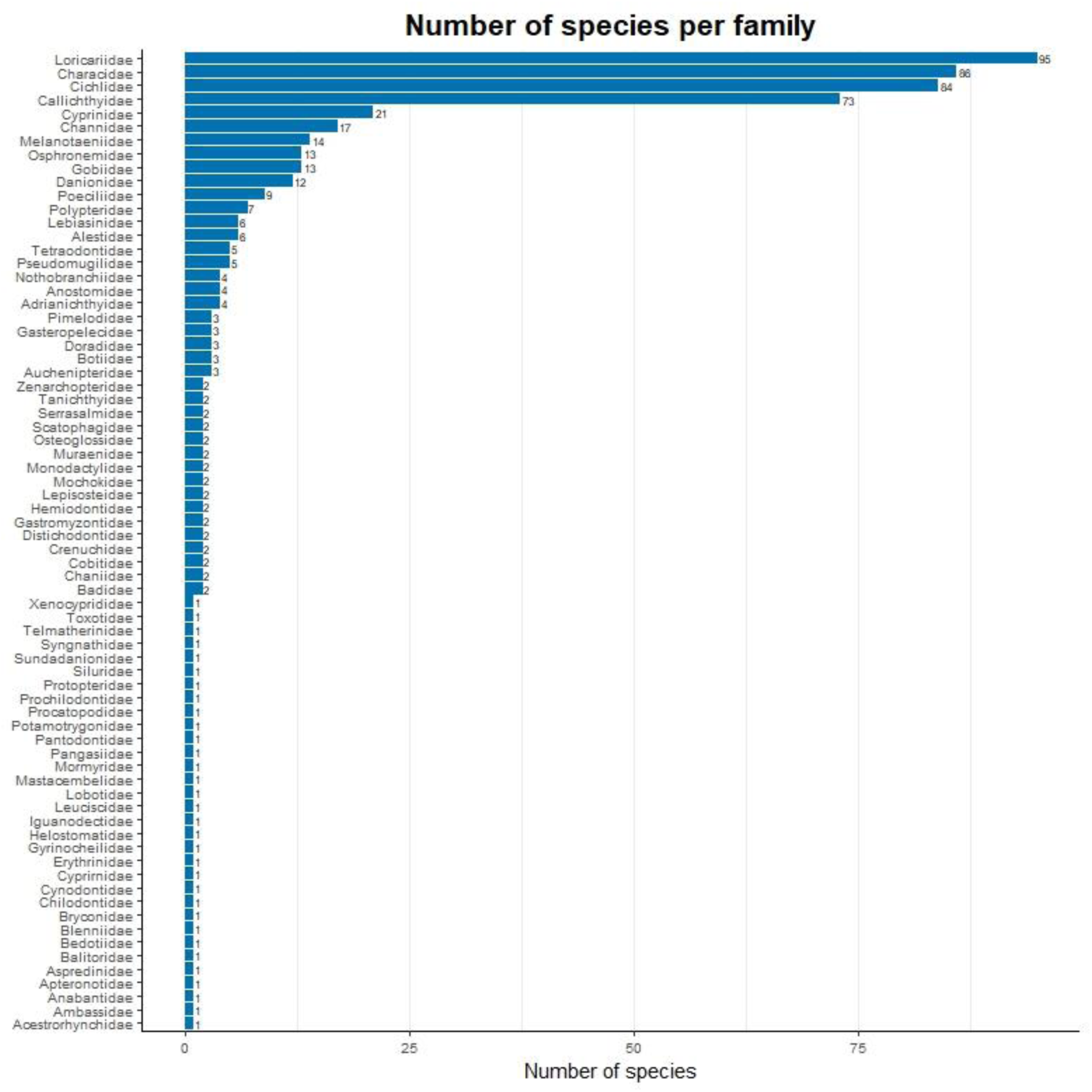
The number of species in different fish families recorded for sale in a pet market in Hong Kong.

**Fig. 3:**
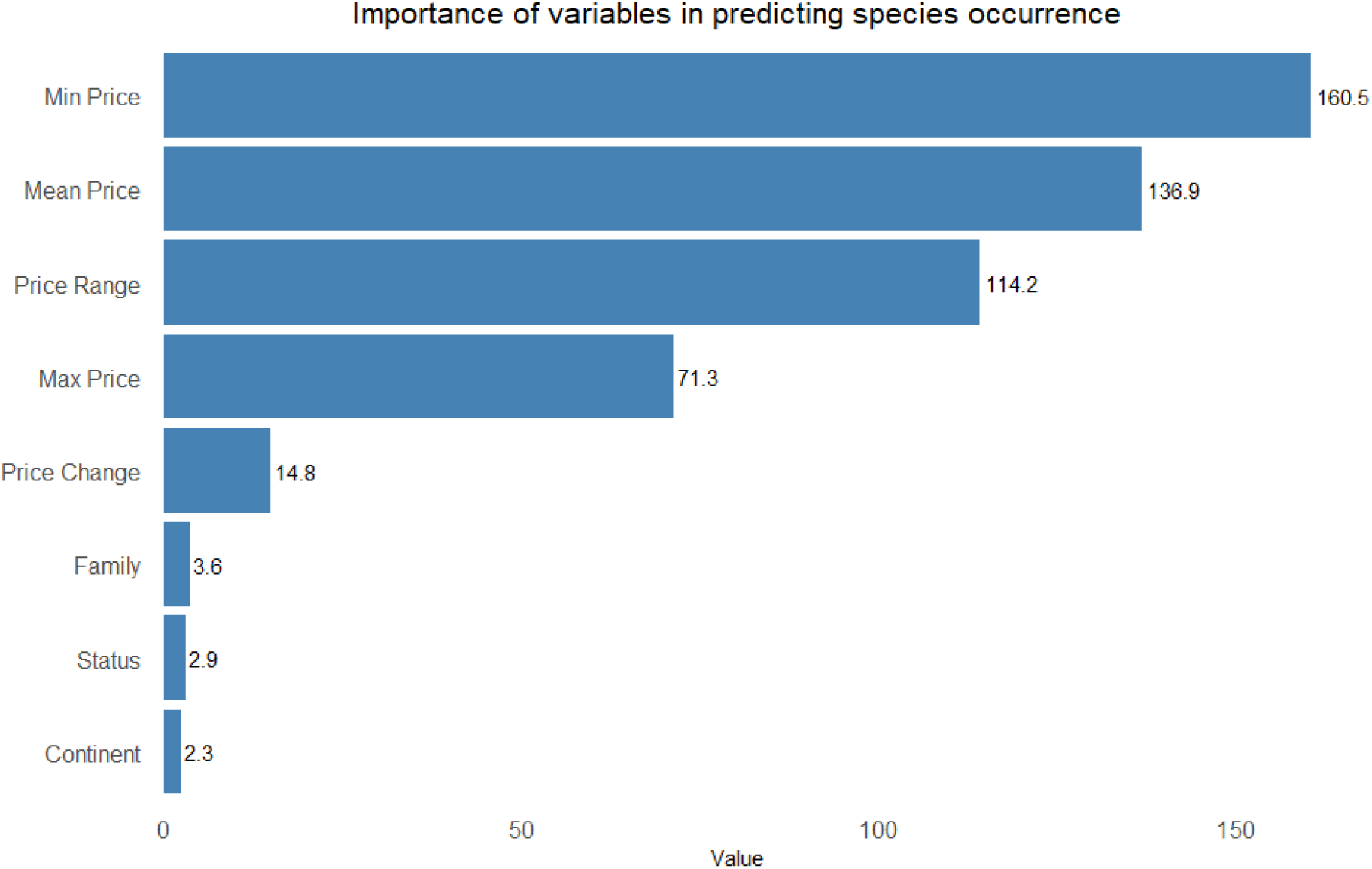
The importance of different variables in predicting species occurrence of freshwater fish sold in a pet market in Hong Kong using random forest analysis.

