## Supplementary Material for "Highlighting the species diversity, pricing trends, and conservation concerns in a major ornamental fish hub"

**Contains:**

**3 supplementary Figures (1 to 3)**

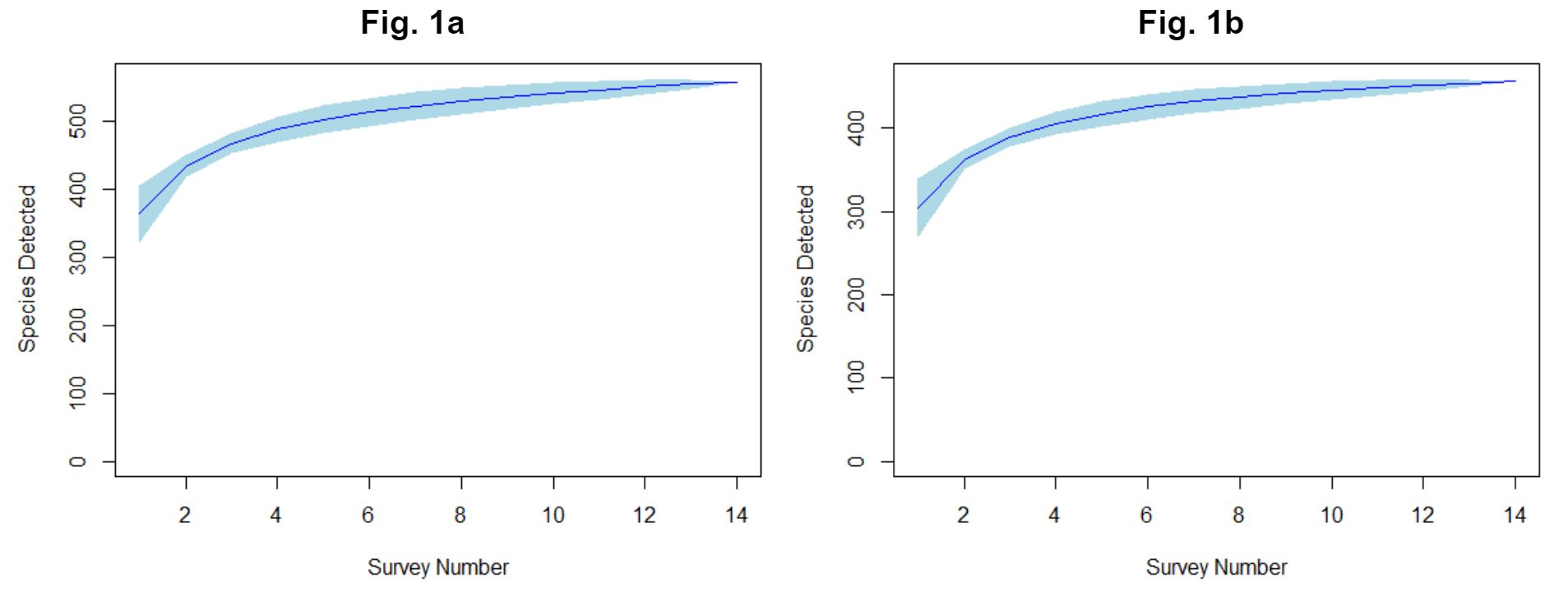

Fig. 1: species accumulation curves from 14 weekly surveys in Gold Fish Street, Mong Kok, Hong Kong using (a) all fishes, and (b) only scientifically recognized fishes. The continuous lines represent the mean and the shaded areas the 95% confidence interval.

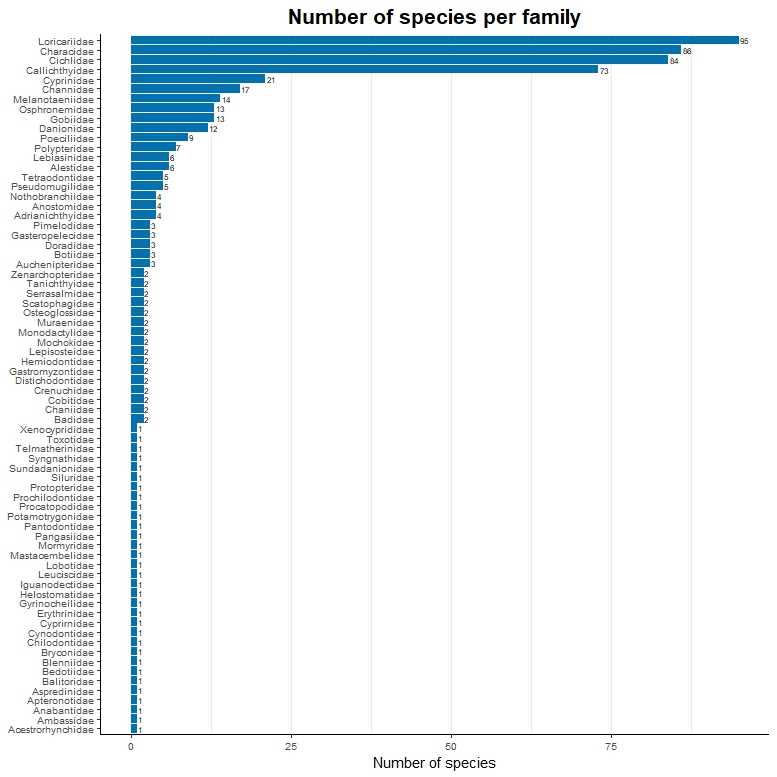

Fig. 2: The number of species in different fish families recorded for sale in a pet market in Hong Kong.

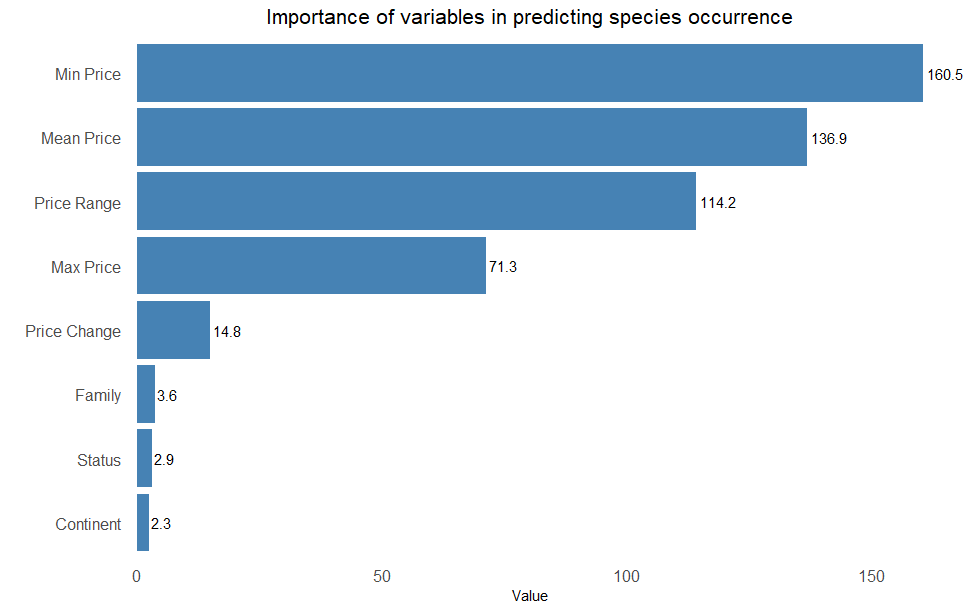

Fig. 3: The importance of different variables in predicting species occurrence of freshwater fish sold in a pet market in Hong Kong using random forest analysis.
